# Whole genome analysis of local Kenyan and global sequences unravels the epidemiological and molecular evolutionary dynamics of RSVgenotype ON1 strains

**DOI:** 10.1101/309187

**Authors:** JR Otieno, EM Kamau, JW Oketch, JM Ngoi, AM Gichuki, Š Binter, GP Otieno, M Ngama, CN Agoti, PA Cane, P Kellam, M Cotten, P Lemey, DJ Nokes

## Abstract

The respiratory syncytial virus (RSV) group A variant with the 72-nucleotide duplication in the G gene, genotype ON1, was first detected in Kilifi in 2012 and has almost completely replaced previously circulating genotype GA2 strains. This replacement suggests some fitness advantage of ON1 over the GA2 viruses, and might be accompanied by important genomic substitutions in ON1 viruses. Close observation of such a new virus introduction over time provides an opportunity to better understand the transmission and evolutionary dynamics of the pathogen. We have generated and analyzed 184 RSV-A whole genome sequences (WGS) from Kilifi (Kenya) collected between 2011 and 2016, the first ON1 genomes from Africa and the largest collection globally from a single location. Phylogenetic analysis indicates that RSV-A transmission into this coastal Kenya location is characterized by multiple introductions of viral lineages from diverse origins but with varied success in local transmission. We identify signature amino acid substitutions between ON1 and GA2 viruses within genes encoding the surface proteins (G, F), polymerase (L) and matrix M2-1 proteins, some of which were identified as positively selected, and thereby provide an enhanced picture of RSV-A diversity. Furthermore, five of the eleven RSV open reading frames (ORF) (i.e. G, F, L, N and P), analyzed separately, formed distinct phylogenetic clusters for the two genotypes. This might suggest that coding regions outside of the most frequently studied G ORF play a role in the adaptation of RSV to host populations with the alternative possibility that some of the substitutions are nothing more than genetic hitchhikers. Our analysis provides insight into the epidemiological processes that define RSV spread, highlights the genetic substitutions that characterize emerging strains, and demonstrates the utility of large-scale WGS in molecular epidemiological studies.

**Author summary:** Respiratory syncytial virus (RSV) is the leading viral cause of severe pneumonia and bronchiolitis among infants and children globally. No vaccine exists to date. The high genetic variability of this RNA virus, characterized by group (A or B), genotype (within group) and variant (within genotype) replacement in populations, may pose a challenge to effective vaccine design by enabling immune response escape. To date most sequence data exists for the highly variable G gene encoding the RSV attachment protein, and there is little globally-sampled RSV genomic data to provide a fine resolution of the epidemiology and evolutionary dynamics of the pathogen. Here we use long-term RSV surveillance in coastal Kenya to track the introduction, spread and evolution of a new RSV genotype known as ON1 (having a 72-nucleotide duplication in the G gene). We present a set of 184 RSV-A whole genomes, including 176 of RSV ON1 (the first from Africa), describe patterns of local ON1 spread and show genome-wide changes between the two major RSV-A genotypes that may define the pathogen’s adaptation to the host. These findings have implications for vaccine design and improved understanding of RSV epidemiology and evolution.

## Introduction

Respiratory syncytial virus (RSV) is the leading viral cause of severe pneumonia and bronchiolitis among infants and children globally (1,2). Individuals remain susceptible to RSV upper respiratory tract reinfection throughout life even though they develop immune responses following primary and secondary RSV infections in childhood (3). No licensed RSV vaccine exists, partly due to the antigenic variability in the virus (4).

The single stranded, negative sense RSV genome encodes 11 proteins of which the attachment glycoprotein (G) is the most variable and a key player of adaptive evolution of the virus (5). The rate of nucleotide substitution for the G gene encoding the attachment protein has been estimated to be 1.83 × 10^−3^ and 1.95 × 10^−3^ nucleotide (nt) substitutions/site/year for group A and B, respectively, with some variation dependent on the timescale of observation (6,7). There is evidence of immune driven selection of the G gene (4,8). Although at a lower rate of evolution than for the G gene, there is significant ongoing accumulation of substitutions across the whole genome, again dependent upon the timescale of observation (9,10). At present, there is limited analysis of the selective forces acting on genes other than for the G gene as a result of paucity of whole genome sequences (WGS), particularly from one location over a period of time spanning multiple seasons (11,12). Therefore, it is not apparent whether there are genetic signatures across the rest of the genome that might additionally inform on the adaptive mechanisms of RSV viruses following introduction into communities.

RSV is classified into two Groups, RSV-A and RSV-B (13), differing antigenically (14), with each group further characterized into genotypes (with genotype defined as a cluster of viruses each of which has greater genetic distance from viruses of any other genotype compared to that between viruses of the most diverse genotype (15,16)). A genotype can be further divided into (i) imported variants which show greater genetic difference than expected from *in situ* diversification (17,18), and (ii) local variants arising from recent introduction which subsequently diversify *in situ* (without time for purifying selection from, for example inter-epidemic bottlenecks) (10). Studies from our group have shown that within RSV epidemics, there is co-circulation of RSV viruses belonging to different groups, genotypes and variants both imported and local, (10,17,18), with the latter not clearly distinguished through partial G gene sequencing. Consequently, full genome sequencing offers the opportunity to differentiate introduced from persistent RSV viruses within a given location.

Two recent RSV genotypes with large duplications within the G glycoprotein, BA and ON1, have been detected globally. The RSV-B BA genotype is characterized by a 60-nucleotide (nt) duplication while the RSV-A ON1 genotype is characterized by a 72-nucleotide duplication. Initially detected in Buenos Aires Argentina in 1999, the BA genotype subsequently spread rapidly throughout the world becoming the predominant group B genotype and replacing all previous circulating RSV-B genotypes in certain regions (19,20). The ON1 genotype was first detected in 2010 in Ontario Canada, a decade after BA, and has also spread globally (21–29). Of interest is what could be driving the apparent fitness advantage of these emergent genotypes over the preceding genotypes (30), and whether such insights could be mined from whole genome sequences.

In this study, we sought to gain a deeper understanding of the epidemiological and evolutionary dynamics of RSV viral populations through extensive whole genome sequencing and analysis of samples collected as part of on-going surveillance studies of respiratory viruses within Kilifi, Coastal Kenya (2011-2016). This was done by monitoring the unique genotype ON1 72-nucleotide duplication tag whose temporal progression can be directly followed from when the ON1 viruses first entered the ON1 ‘naïve’ Kilifi population. This WGS analysis advances previous work on the patterns of introduction and persistence of the ON1 variant within this community that utilized partial G gene sequences (31,32), and provides a higher resolution of the RSV genetic structure, spread and identification of variation that may be associated with molecular adaptation and apparent fitness advantages.

## Materials and Methods

### Study population

This study is part of ongoing surveillance of respiratory viruses within Kilifi County, coastal Kenya, and across the country that is aimed at understanding the epidemiology and disease burden of respiratory viruses in this region (33). Two sets of samples were used in the current analysis; (i) samples collected from children (under 5 years of age) admitted to the Kilifi County Hospital (KCH) presenting with syndromically defined severe or very severe pneumonia (32,33), and (ii) samples collected from patients of all ages presenting at health facilities within the Kilifi Health and Demographic Surveillance System (KHDSS) (34) with acute respiratory illnesses (ARI).

### RNA extraction and RT-PCR

All KCH specimens had previously been screened for RSV, RSV group and RSV-A genotype status (32), while the KHDSS samples were screened afresh. The criterion for proceeding to WGS was a sample real-time PCR cycle threshold (Ct) value < 30 based on the success rate from previous experience (9), with the exception of four test samples that were PCR negative or had Ct>30. Viral RNA was extracted using QIAamp Viral RNA Mini Kit (QIAGEN). Reverse transcription (RT) of RNA molecules and polymerase chain reaction (PCR) amplification were performed with a six-amplicon, six-reaction strategy (9), or using a 6 or 14-amplicon strategy (unpublished) split into two reactions of three and seven amplicons, respectively for each, *Figure 1A*. Amplification success was confirmed by observing the expected PCR product size (1200-1500 bp) on 0.6% agarose gels. For the successfully amplified samples, all the six or two reactions per sample were pooled and purified for Illumina library preparation.

**Figure 1:**
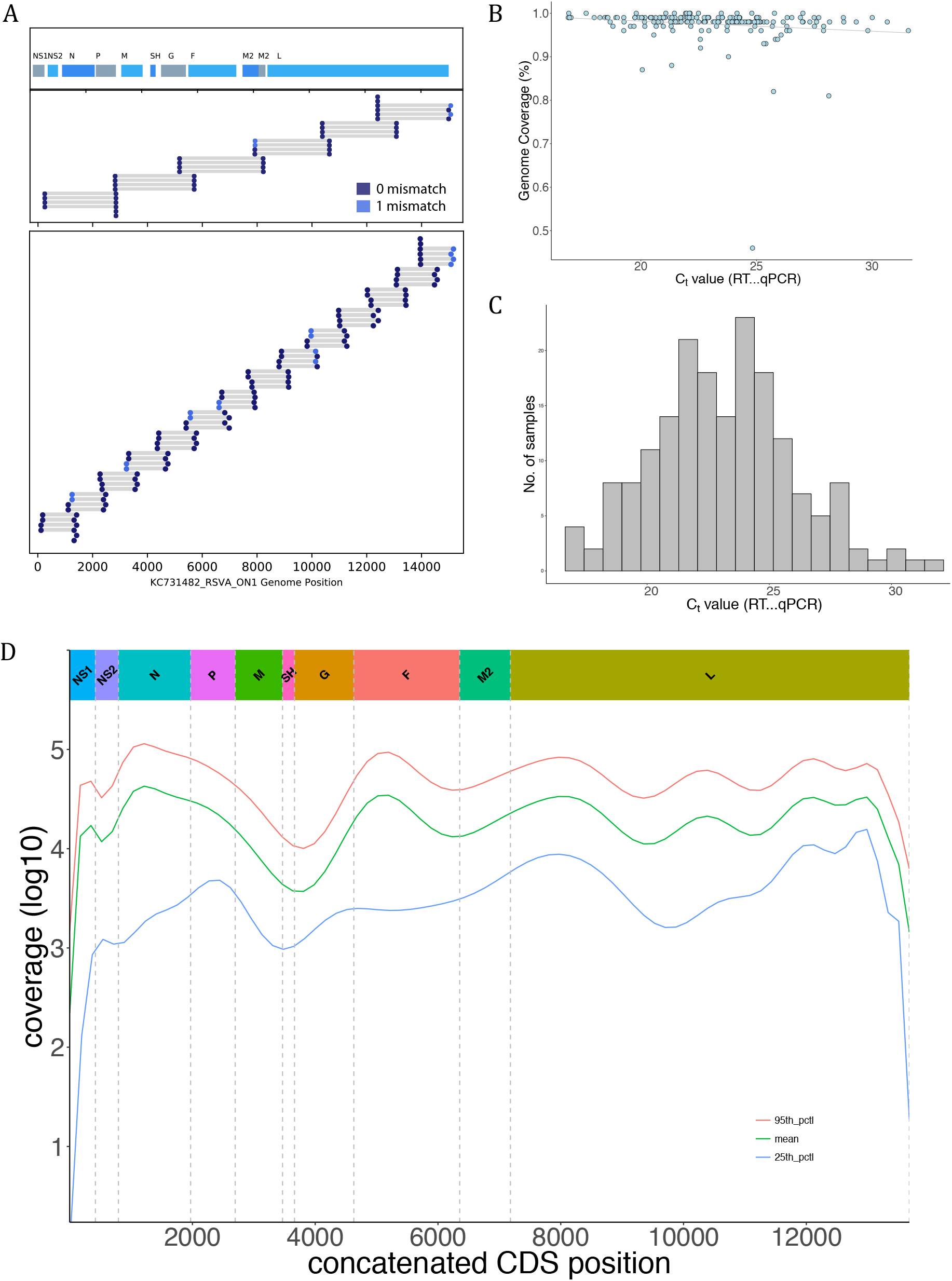
Sample sequencing and genome details. The two RSV-A whole genome amplification strategies used in this study are shown in (A), i.e. six and fourteen amplicons. For each panel the positions of primer targets for each amplicon are indicated. The locations of the 11 RSV ORFs are indicated on top of panel 1. (B) The proportion of RSV genome length sequence recovered (using KC731482 as the reference) was plotted as a function of sample’s diagnostic real-time PCR Ct value. (C) The distribution of the diagnostic real-time PCR Ct values for the samples reported here. (D) Shows the log values of the sequencing depth (see Methods) at each position of the genome assemblies along the concatenated RSV ORFs (i.e. excluding the intergenic regions).

### Illumina library construction and sequencing

The purified PCR products were quantified using Qubit fluorimeter 2.0 (Life Technologies) and normalized to 0.2 ng/μL. The normalized DNA was tagmented (a process of fragmentation and tagging) using the Nextera XT (Illumina) library prep kit as per the manufacturer’s instructions. Indices were ligated to the tagmented DNA using the Nextera XT index kit (Illumina). The barcoded libraries were then purified using 0.65X Ampure Xp beads. Library quality control was carried out using the Agilent high sensitivity DNA kit on the Agilent 2100 Bioanalyzer (Agilent) to confirm the expected size distributions and library quality. Each library was quantified using the Qubit fluorimeter 2.0 (Life Technologies), after which the libraries were then normalized and pooled at equimolar concentrations based on the Qubit results. The pooled libraries were sequenced on either (i) Illumina HiSeq system using 2 × 250 bp PE sequencing at the Wellcome Trust Sanger Institute (UK), or (ii) Illumina MiSeq using 2 × 250 bp PE sequencing at the KEMRI-Wellcome Trust Research Programme (Kilifi, Kenya).

To determine the proportion of RSV and non-RSV reads in the samples used here, Kraken v0.10.6 (35) was used with a pre-built Kraken database provided by viral-ngs (36,37) (downloaded in December, 2015; https://storage.googleapis.com/sabeti-public/meta_dbs/kraken_ercc_db_20160718.tar.gz). A preliminary quality check of the sequence reads was done using fastqc (38) with the output per batch aggregated and visualized by multiqc (39).

### Depletion of human reads

Prior to deposition of the raw short reads into NCBI short read archive (SRA), datasets were depleted of human reads. The raw reads were mapped onto the human reference genome hg19 using bowtie2 (40) while samtools (41) was used to filter, sort and recover the unmapped (non-human) reads. The final reads are available in the NCBI BioProject database under the study accession PRJNA438443.

### Genome assembly and coverage

Sequence reads were taxonomically filtered within the viral-ngs pipeline using an RSV genotype ON1 reference, KC731482. The RSV reads were then used to generate consensus genome assemblies using viral-ngs versions 1.18.0 and 1.19.0 (36,37) and/or SPAdes version 3.10.1 (42), selecting the most complete assembly from either assemblers. In addition, the available Sanger G-gene sequences (31,32) for these same samples were used to confirm agreement with the WGS assemblies. The genomes generated in this study are available in GenBank under accession numbers MH181878 - MH182061. The genomes were aligned using MAFFT alignment software v7.305 (43) using the parameters ‘--localpair --maxiterate *1000*’

To calculate and visualize depth of coverage, sample raw reads were mapped onto individual assemblies with BWA (44), samtools (41) used to sort and index the aligned bam files, and finally bedtools (45) used to generate the coverage depth statistics. Plotting of the depth of coverage was done in R (46) in the RStudio (47).

### Global comparison dataset

All complete and partial genome sequences available in GenBank Nucleotide database (https://www.ncbi.nlm.nih.gov/genbank/) as on 19/09/2017 were added to a global RSV-A genotype ON1 genomic and G-gene dataset. To prepare the global ON1 dataset, we downloaded all RSV sequences from GenBank (search terms: respiratory syncytial virus), created a local blast database in Geneious (48), and performed a local blast search using the 144 nucleotide sequence region of the ON1 genotype. To remove duplicates, the sequences were binned by country of sample collection, filtered of duplicates and then re-collated into a single dataset. For the global G-gene dataset of 1,167 sequences, the sequence length ranged from 238-690bp. The final alignment of 344 ON1 genome sequences comprised the sequences reported in this study (*n*=154) and additional publicly available GenBank ON1 sequences (*n*=190). In addition to the ON1 genomes, we generated 30 genotype GA2 genome sequences from Kilifi. The alignments were inspected in AliView (49) and edited manually removing unexpected spurious frame-shift indels (largely homopolymeric and most likely sequencing errors).

### Maximum likelihood phylogenetic analyses and root-to-tip regression

Separate Maximum-Likelihood (ML) phylogenetic trees were generated using multiple sequence alignments of the three datasets, i.e. Kilifi WGS, and global G-gene and WGS datasets. The ML trees were inferred using both PhyML and RaxML, with each optimizing various parts of the tree generation process (i.e. borrowing strengths of both approaches), using the script generated and deposited by Andrew Rambaut at (https://github.com/ebov/space-time/tree/master/Data/phyml_raxml_ML.sh). The GTR+G model was used after determination as the best substitution model by IQ-TREE v.1.4.2 (50).

To determine presence of temporal signal (‘clockiness’) in our datasets, we used TempEst v1.5 (51) to explore the relationship between root-to-tip divergence and sample dates. The data were exported to R (46) to perform a regression with the ‘lm’ function.

### Estimating number local variant introductions

To differentiate between local variants arising from a recent introduction and imported variants with greater genetic differences than is expected from local diversification, we used a pragmatic criterion previously described by *Agoti et al*. in (17). Briefly, a variant is a virus (or a group of viruses) within a genotype that possesses ≥*x* nucleotide differences compared to other viruses. This *x* nucleotide differences is a product of the length of the genomic region analyzed, estimated substitution rate for that region, and time. This analysis was done using usearch v8.1.1861 (29).

### Protein substitution and selection analysis

Using the aligned Kilifi (ON1 and GA2) genome dataset, patterns of change in nucleotides (single nucleotide polymorphisms or SNPs) and amino acids were sought using Geneious v11.1.2 (48) and BioEdit 7.2.5 (52), respectively. Potential positively selected and co-evolving sites within the coding regions were identified using HyPhy (53) and phyphy (54). SNPs were called from both the complete dataset and from an alignment of the consensus sequences from GA2 and ON1, whereby a consensus nucleotide was determined as the majority base at a given position. For the positive selection analysis, two strategies were used; gene-wide selection detection [BUSTED (55)] and site-specific selection [SLAC, FEL (56), FUBAR (57) and MEME (58)]. Codon positions with a p-value <0.1 for either the SLAC, FEL and MEME models or with a posterior of probability >0.9 for the FUBAR method were considered to be under positive selection.

### Bayesian phylogenetics

To infer time-structured phylogenies, Bayesian phylogenetic analyses were performed using BEAST v.1.8.4 (59). Because of sparse data at the 5’ and 3’ termini and in the non-coding regions of the genomic datasets, only the coding sequences (CDS) were used as input. The SRD06 substitution model (60) was used on the CDS and three coalescent tree priors were tested, i.e. a constant-size population, an exponential growth population, and a Bayesian Skyline (61). For each of these tree priors, combinations with the strict clock model and an uncorrelated relaxed clock model with log-normal distribution (UCLN) (62) were tested with the molecular clock rate set to use a non-informative continuous time Markov chain rate reference prior (CTMC) (63). For each of the molecular clock and coalescent model combinations, the analyses were run for 150 million Markov Chain Monte Carlo (MCMC) steps and performed both path-sampling (PS) and stepping-stone (SS) to estimate marginal likelihood (64,65). The best fitting model was a relaxed clock with a Skyline coalescent model, *Supplementary sheet 1*.

BEAST was then run with 300-400 million MCMC steps using the SRD06 substitution model, Skyline tree prior, and relaxed clock model to estimate Bayesian phylogenies. For the time to the most recent common ancestor (TMRCA) estimates, the same substitution model and tree prior were used as above but with a strict clock model. For the global G-gene dataset, BEAST was run with 400 million MCMC steps using the HKY substitution model, Skyline tree prior, and a relaxed clock model. We used Tracer v1.6 to check for convergence of MCMC chains and to summarize substitution rates. Maximum clade credibility (MCC) trees were identified using TreeAnnotator v1.8.4 after removal of 10% burn-in and then visualized in FigTree v1.4.3.

### Principal component analysis

To check on any clustering and stratification patterns, principal component analysis (PCA) was performed using the R package FactoMineR (66). The input data were a matrix of pairwise distances from genome sequence alignment using the “N” model of DNA evolution, i.e. the proportion or the number of sites that differ between each pair of sequences. Each genome on the PCA plot was annotated by the continent of sample origin.

## Results

### Genome sequencing and assemblies

A total of 184 RSV-A genomes were generated in this study, comprising genotypes ON1 (*n*=154) and GA2 (*n*=30), collected between February 2012 and April 2016; *Supplementary sheet 2*. This dataset included 176 genomes from inpatients at KCH and 8 genomes from peripheral health centres within the KHDSS. Between 0.2 to 4.3 million short reads were available per sample of which RSV specific reads ranged between 0.001 to 3.9 million reads. The genome assemblies had a median length of 15,054 nucleotides (range: 13,966-15,322) and mean depth of base coverage per genome ranging from 39 to 66457.

Whereas the samples for WGS were generally of high viral content (lower Ct value), it is apparent there was reduced genome yield (proportion of genome assembled) from samples with lower viral loads (i.e. higher Ct values); *Figure 1B*. However, since the samples collected at the hospital are from children presenting with severe or very severe pneumonia cases they generally have high viral loads as shown in *Figure 1C*. The median fraction of the genome with unambiguous base calls was 98% with reference length from KC731482. Read coverage across the genomes was non-uniform, *Figure 1D*, suggesting varied PCR amplification efficiency among primer pair combinations combined with increased sequencing yield from the ends of the amplicons.

### Bayesian reconstruction of ON1 epidemiological and evolutionary history

The global ON1 whole-genome MCC phylogenetic tree, *Figure 2A*, shows evolutionary relationship among ON1 viruses from five sampled continents. The TMRCA of the ON1 strains from the most recent tip (7 April 2016) was estimated to be 11.07 years [95% HPD: 9.85-12.31], resulting in an estimated ON1 emergence date of between December 2003 and June 2006. This estimated date of emergence is earlier than a previous estimate (2008-2009) using the G-gene alone (29), but such a difference could ideally be a reflection of the different datasets (by geography and sampling dates). *Comas-Garcia et al*. have reported the earliest ON1 strain identified to date in November 2009 from central Mexico (67), and from our estimates this suggests a period of 3-6 years of circulation of this virus before first detection. The genome-wide substitution rate for the ON1 viruses was estimated at 5.97 × 10^−4^ nucleotide substitutions per site per year [95% HPD: 5.42-6.58], similar to previous estimates for RSV group A full length sequences sampled over several epidemics (9,12) but slower than estimates from both samples collected from a household study over a single epidemic within the same location and using a global ON1 G-gene dataset (10,29). Across the genome, estimates of evolutionary rates for individual ON1 open reading frames (ORFs) varied, *Figure 2B*, with the mean substitution rate highest in the G-gene, lowest in NS1, and moderate (with tight 95% HPD intervals) for the whole genome.

**Figure 2:**
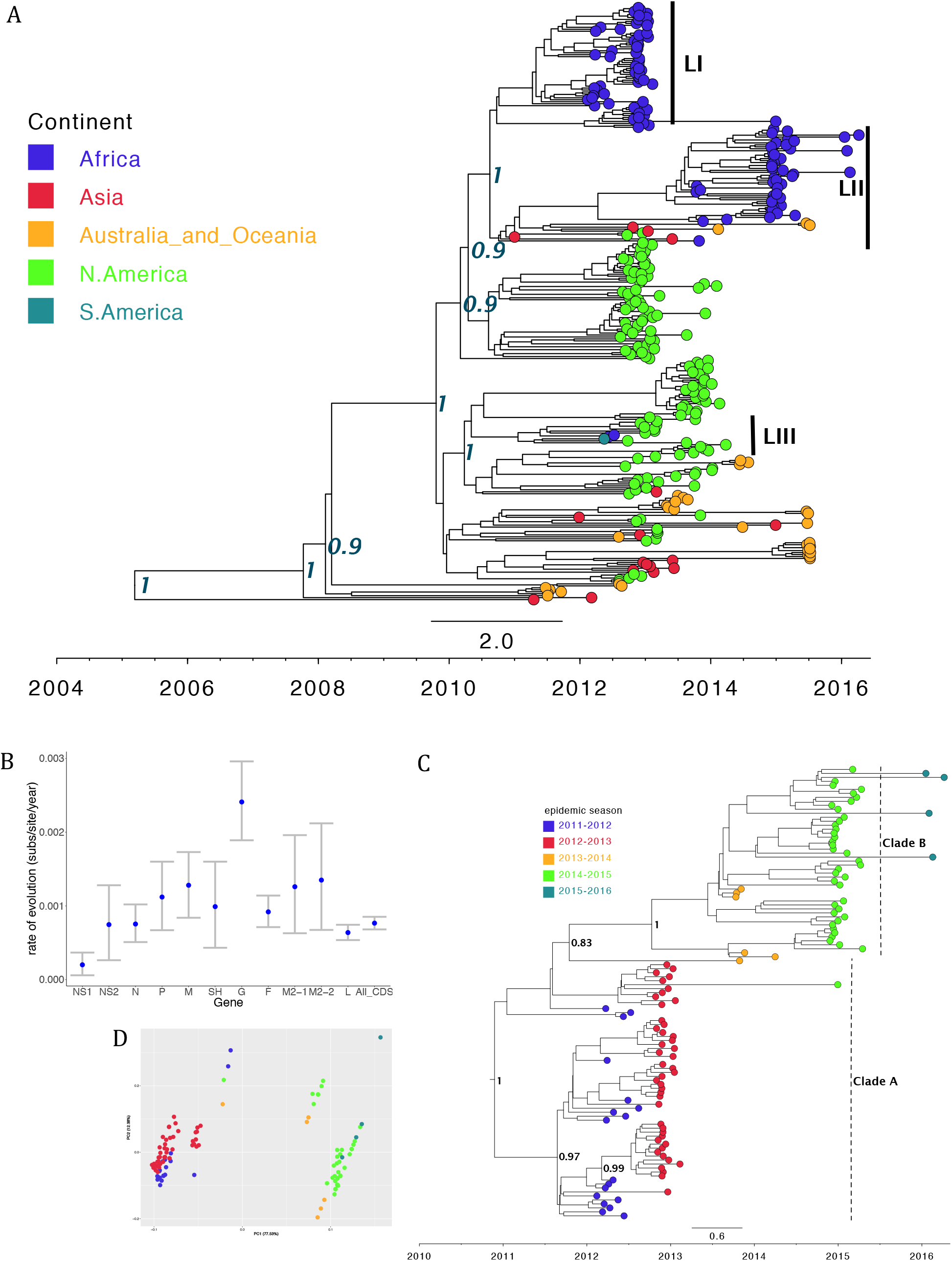
Global and local ON1 MCC trees and PCA. (A) Maximum clade credibility tree inferred from 344 global full genome sequences (see Methods), annotated with the identified Kilifi lineages (I-III) and the tips colour coded with the continent of sample collection. Node labels are posterior probabilities indicating support for the selected nodes. (B) shows the evolutionary rate estimates for the different genotype ON1 ORFs. (C) is an MCC tree inferred from 154 ON1 genomes from Kilifi annotated with identified clades A and B, and the tips colour coded with the epidemic season. (D) is a PCA analysis (see Methods) of the same dataset as (C) and similarly annotated. Percentage of variance explained by each component is indicated on the axis.

The Kilifi ON1 genomes formed three distinct lineages (labelled LI-LIII) on the global tree in *Figure 2A*. These three ON1 lineages were however placed into two clades within the Kilifi WGS MCC tree, *Figures 2C* whereby lineages LI and LIII were placed in clade A while clade B comprised only lineage LII sequences. The two clades were temporal with clade A mostly comprising sequences from the 2011-2013 RSV epidemic period and clade B comprising sequences from the epidemic period 2013-2016. These clade and temporal patterns are further highlighted by the PCA analysis in *Figure 2D*. Based on the phylogenetic placement of the Kilifi ON1 lineages on the global tree, we estimate that there could have been at least three separate introductions of ON1 viruses into Kilifi. For lineage LIII, we sampled only 4 cases, and this is consistent with limited local transmission. In addition, all the eight outpatient ON1 viruses collected outside KCH were placed within lineage LII and were interspersed with viruses sampled from inpatient admissions at KCH implying that our sampling at the hospital might be well representative of the KHDSS community.

Using the global whole genome ON1 substitution rate estimate above, the Kilifi ON1 genomes dataset (length 15,404 bp) and a pragmatic criterion previously described by *Agoti et al*. in (17) to differentiate between local and imported variants, we estimated that there were up to 73 ON1 introductions into Kilifi. Even when we used the higher substitution rate previously estimated from ON1 partial G-gene sequences by *Duvvuri et al* in (29), i.e. 4.10 × 10^−3^ substitutions/site/year which translates to a difference of at least 63 nucleotides between any two genomes to be classified as separate introductions, this resulted in an estimate of 6 separate introductions. This implies that multiple seeding introductions of viruses within lineages LI and LII may have been required to sustain their local transmission.

### Global ON1 spatiotemporal dynamics

As there are far more partial G gene sequences than full genomes, we explored ON1 spatiotemporal patterns using a set of 1,167 global G gene sequences. The global G gene MCC tree is shown in *Figure 3* with the corresponding sampling locations in *Supplementary figure 1*. Although they do not correspond to well-supported monophyletic clades, we classify the clusters in C1, C2 and C3 for convenience. While there was neither a single cluster that was comprised solely of viruses from a specific continent nor a continent whose viruses were only found within a single cluster, there was predominance of African/European viruses in cluster C1, Asian in C2 and European/Asian in C3, suggesting both intra and inter-continental circulation patterns. The majority of the Kilifi ON1 lineages (LI and LII in *Figure 2A*) were found in cluster C1, suggesting perhaps a predominantly European source of RSV introductions into Kilifi, while the lineage LIII viruses were found in cluster C3. All the African viruses in cluster C1 were from Kilifi while all the Nigerian and all the South African viruses were only found in clusters C2 and C3 respectively. Further, the viruses closely related to the ON1 lineage (LIII) with limited transmission in Kilifi described above were frequently isolated in other locations (cluster C3).

**Figure 3:**
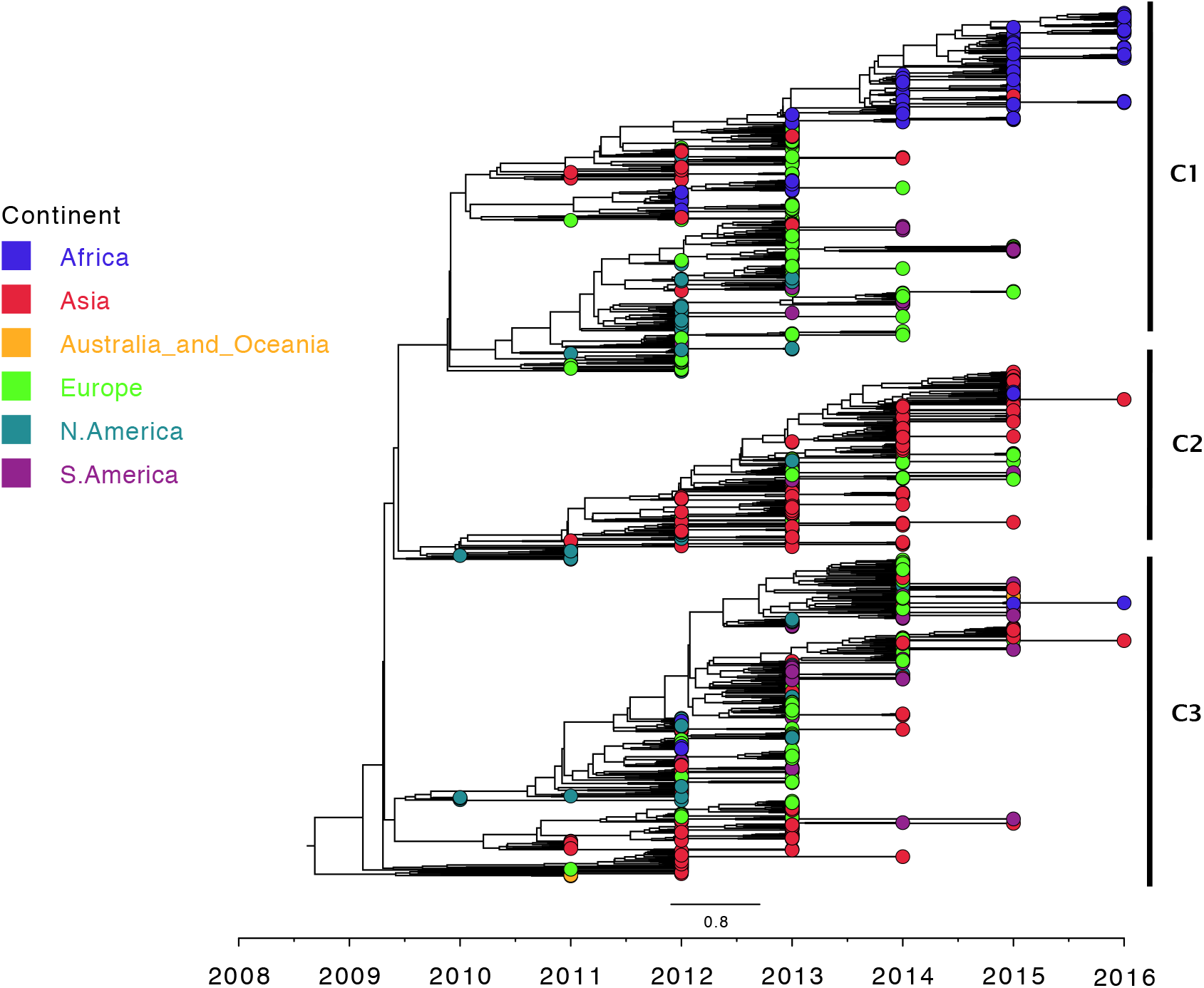
Global ON1 G-gene MCC phylogenetic tree. A maximum clade credibility tree inferred from 1,167 partial ON1 G gene global sequences with the tips colour coded with the source continent.

### Genomic diversity of Kilifi RSV-A viruses

Pairwise intra-genotypic genetic diversity analysis of the GA2 and ON1 genomes from Kilifi, *Figure 4A* and *4B*, show unimodal and bimodal distributions respectively consistent with two genetically distinct circulating strains of the ON1 viruses. *Figure 4C* shows an entropy plot with protein substitution density based on amino acid polymorphisms for a concatenated set of all RSV proteins from both Kilifi genotype ON1 and GA2 viruses. Analyzing for substitutions across the genomes, we identified a total of 746 single nucleotide polymorphisms (SNPs) with frequencies of >1% (in the set of 184 genomes). Of these SNPs, the majority (589, 78.9%) were found within coding sequences/regions (CDS) with only 145/589 (24.6%) of these coding mutations resulting in non-synonymous changes, *Supplementary sheet 3*. The three CDSs with the most substitutions were the polymerase L (39.6%), the glycoprotein G (14.8%) and the fusion F (14.6%). However, most of the non-synonymous changes occurred within G, SH and M2-2.

**Figure 4:**
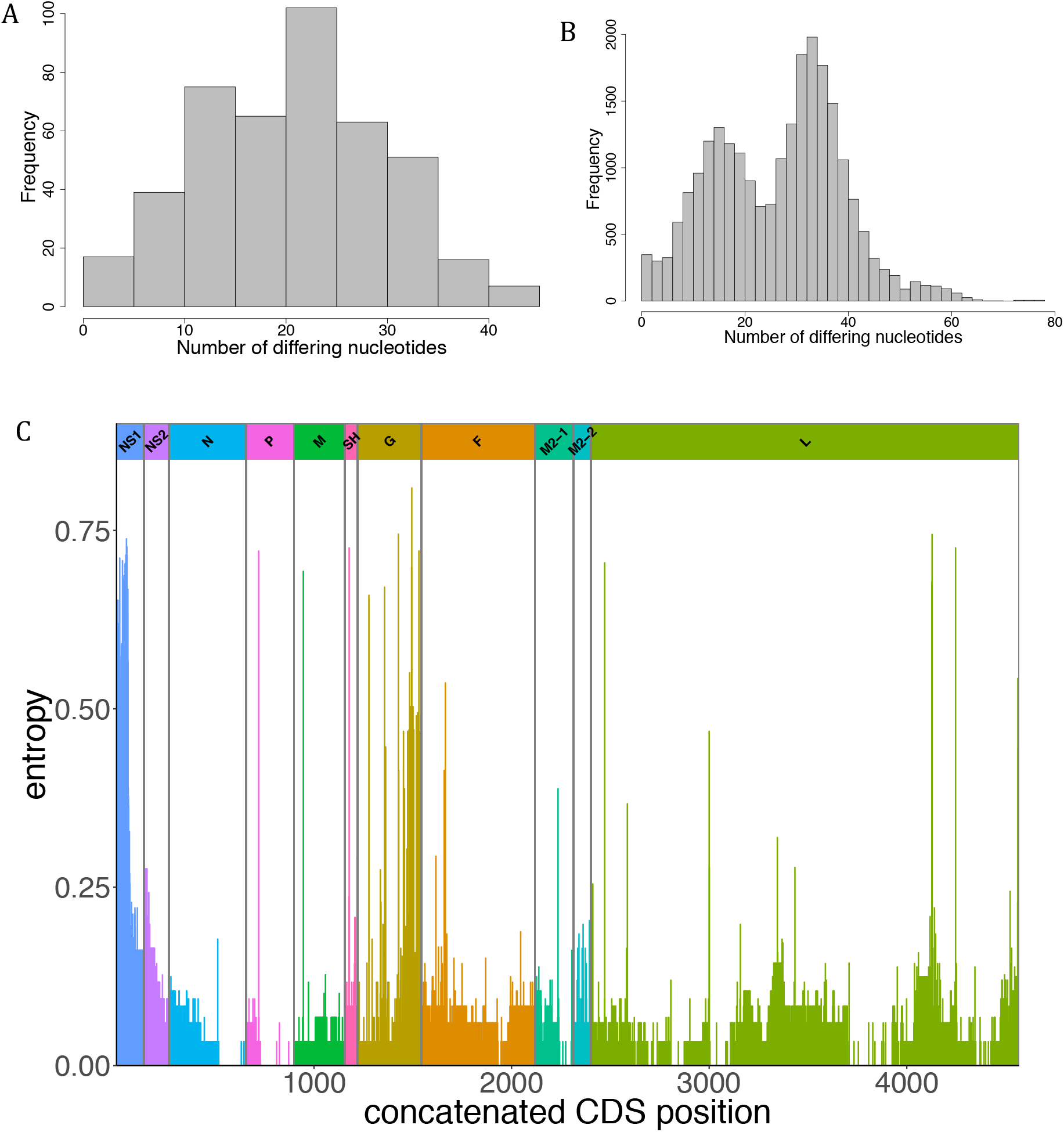
Pairwise genomic distances and genome-wide amino acid variation. The distribution of pairwise genetic distances between genotype GA2 and ON1 genome sequences are shown in (A) and (B), respectively. (C) is an entropy plot showing amino acid variation along the concatenated Kilifi RSV-A genomes.

### Is the 72-nucleotide duplication a red-herring or masking the complete genomic identity of ON1 viruses?

The currently known or *de facto* distinguishing feature of the ON1 from GA2 strains is the 72-nucleotide duplication within the G gene. It has been shown from phylogenetic analysis of the G-gene that RSV group A genotypes form distinct clusters (16). However, it has not been investigated if the distinct clustering is replicated in the other genes especially for the closely related genotypes GA2 and ON1 viruses. Through an exploratory root-to-tip regression analysis of ORF specific ML trees, we confirmed that all but the NS1, NS2 and SH proteins had good temporal signals to proceed with this analysis, *Supplementary figure 2*. Our observations using the Kilifi GA2 and ON1 WGS dataset indicate that this phylogenetic divergence is present in the concatenated set of all the 11 RSV-A ORFs (*Figure 5A*) and five individual coding regions (G, F, L, N and P), *Supplementary figure 3*. However, node posterior support for divergence between GA2 and ON1 in the N and P proteins was quite low (50-70%) despite observation of distinct clusters. The date of the MRCA of the Kilifi ON1 strains was estimated to June 2010 [95% HPD: November 2009-November 2010], implying a lapse of at least one year before initial detection of genotype ON1 in Kilifi in February 2012.

**Figure 5:**
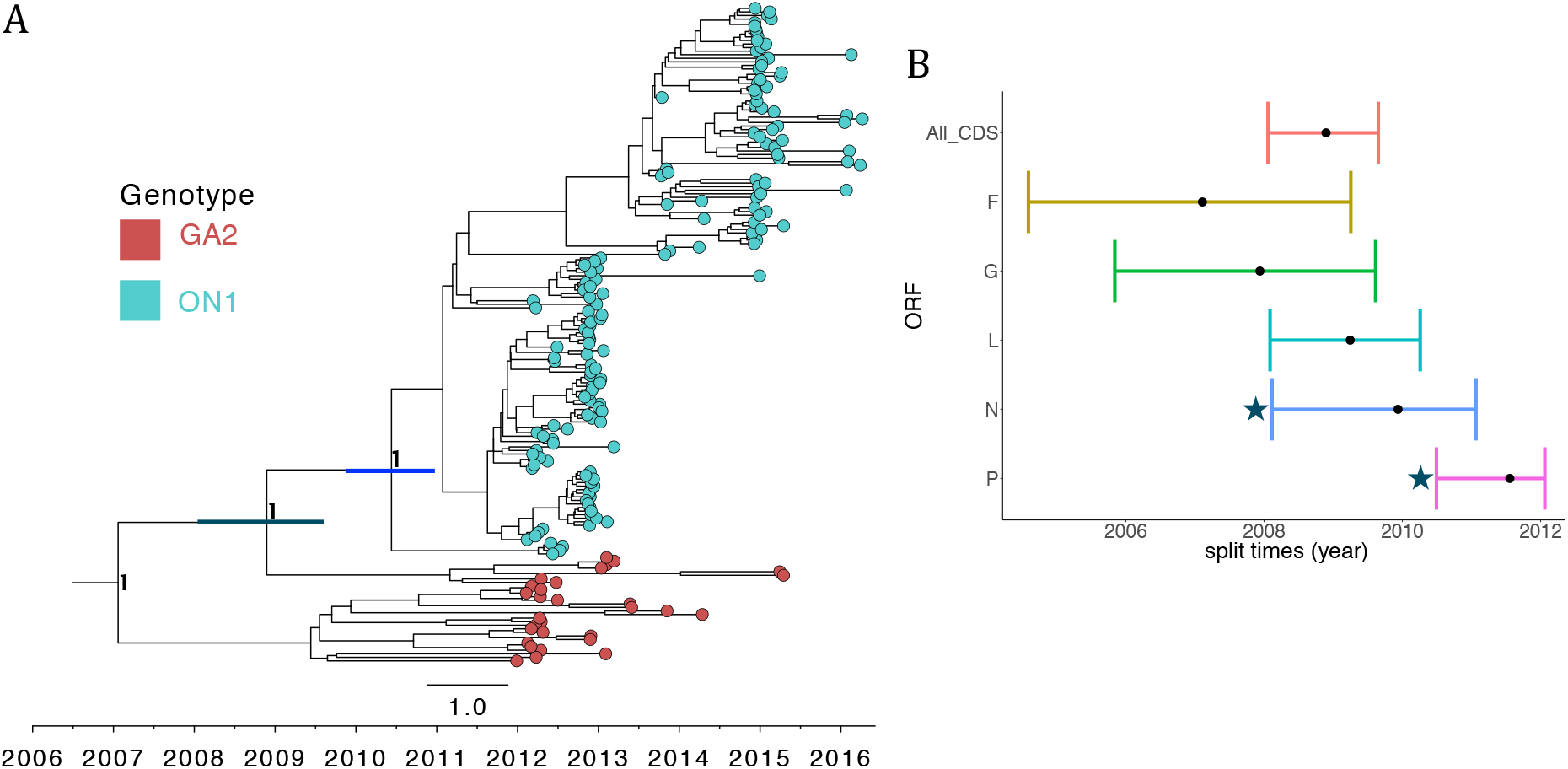
Estimated TMRCA for Kilifi RSV-A viruses and ORFs. (A) Maximum clade credibility tree inferred from 184 RSV-A complete genome sequences (coding regions only) from Kilifi with the tips colour coded by genotype, i.e. ON1 (cyan) and GA2 (red). The two node bars indicate the 95% HPD interval for the TMRCA for the Kilifi GA2 and ON1 viruses (grey), and Kilifi ON1 strains (blue). Node labels are posterior probabilities indicating support for the selected nodes. (B) shows the TMRCA (with 95% HPD interval) of the different ORFs of the RSV-A genotype GA2 and ON1 viruses. The (*) indicates node posterior support of <0.9 for the split between GA2 and ON1 in the N and P ORFs.

Based on the results above indicating evolutionary divergence across the five ON1 and GA2 proteins, we asked the question; Is the 72-nucleotide duplication the only marker of the ON1 strains or an accompanying mutation? To answer this question, we analyzed a concatenated set of 10 RSV ORFs (excluding the G) whereby we observed distinct and well supported ON1 and GA2 clusters indicating the presence of genetic markers outside the 72-nucleotide duplication and the G ORF that differentiate viruses belonging to these two genotypes. With the possibility of additional substitutions between GA2 and ON1 viruses across the genome and assuming a single point source of ON1 viruses, we hypothesized two likely scenarios (i) a single ON1-GA2 split event in which the founder ON1 virus possessed distinctive substitutions across the five RSV ORFs, or (ii) progressive but rapid accumulation of substitutions between ON1 and GA2 viruses within the five ORFs.

With regard to scenario (ii) above, it would be important to know what could have come first; the 72-nucleotide duplication in the G or the changes in the other ORFs? However, considering that these substitutions could have happened anywhere on the branch between the GA2-ON1 split time and ON1 TMRCA in *Figure 5A*, it is impossible to distinguish the order of such substitutions. This dilemma is confirmed by the overlapping intervals in the divergence timings of the individual ORFs in *Figure 5B*.

### Identification of signature substitutions differentiating ON1 from GA2 viruses

The presence of phylogenetic divergence between five ORFs of ON1 and GA2 viruses above indicates potential SNPs between these two strains. Through a comparative genome-wide scan along the RSV-A coding genome, we sought to pick out SNPs between the consensus sequences of the Kilifi ON1 and GA2 viruses. We identified 66 signature nucleotide substitutions, i.e. where a signature substitution was defined as an SNP differentiating ON1 from GA2 viruses, *Supplementary sheet 4*. While most of these signature substitutions were synonymous, we found 14 non-synonymous substitutions between the ON1 from GA2 viruses, *Table 1*, of which nine substitutions were in the G protein, two each in the F and L proteins, and one in the M2-1 protein. None of these signature substitutions were observed to have an effect on our RSV multiplex PCR diagnostics as they occur outside the target primer binding sites in the N gene. Changes at the codon sites 142 and 237 of the G protein that had these signature substitutions have previously been shown to characterize antibody escape mutants, and were located within strain-specific epitopes (68). The two signature substitutions in the F protein (116 and 122) occur within site p27, which is the most variable antigenic site in the F protein among RSV group A and B genotypes (69). However, determining the effect of these signature AA substitutions on potential fitness and phenotype differences between ON1 and GA2 strains will require targeted functional assays.

**Table 1:** Signature nucleotide and amino acid polymorphisms between genotype ON1 and GA2 viruses

| ORF | <sup>a</sup>ORF Nt Pos. | ORF AA Pos. | Change | AA Change | SNP Type |
| --- | --- | --- | --- | --- | --- |
| G | 424 | 142 | TT -> CA | L -> Q | Substitution |
| G | 622 | 208 | C -> A | L -> I | Transversion |
| G | 695 | 232 | G -> A | G -> E | Transition |
| G | 709 | 237 | A -> G | N -> D | Transition |
| G | 758 | 253 | A -> C | K -> T | Transversion |
| G | 817 | 273 | T -> A | Y -> N | Transversion |
| G | 821 | 274 | C -> T | P -> L | Transition |
| G | 851 | 284 | 72 nt<br>duplication | 24 AA<br>insertion | Deletion |
| G | 929 (GA2: 857) | 310 | C -> T | P -> L | Transition |
| G | 941 (GA2: 869) | 314 | T -> C | L -> P | Transition |
| F | 346 | 116 | A -> G | N -> D | Transition |
| F | 364 | 122 | G -> A | A -> T | Transition |
| M2-1 | 349 | 117 | A -> C | N -> H | Transversion |
| L | 1792 | 598 | C -> T | H -> Y | Transition |
| L | 5175 | 1725 | A -> T | E -> D | Transversion |

ORF=Open Reading Frame, Nt=Nucleotide, AA=Amino Acid, Pos.=Position
<sup>a</sup>Positions are relative to ON1 strains, in which complementary positions in GA2 (without the duplication) within the G protein are shown in brackets.

It should be noted that the signature ON1-GA2 SNPs above were called from consensus genome sequences of each genotype, whereby a consensus nucleotide was a simple majority at a given position. However, five nucleotide positions with the identified signature polymorphisms resulting in non-synonymous substitutions had 100% nucleotide consensus for GA2 and ON1 sequences and are of great interest, i.e. codon positions 232, 253, 274 and 314 in the G protein and 598 in the L protein (respective nucleotide positions in *Table 1*). Additionally, we observed the following substitutions in ON1 genomes (data not shown) that might also be of interest; (i) ON1 viruses undergoing convergent evolution at particular sites by acquiring nt/AA substitutions similar to GA2 viruses in recent epidemics (2014-2016), and (ii) ON1 viruses undergoing further diversification through acquisition of nt and/or AA that is different from GA2 in recent epidemics whereas these two genotypes possessed different or shared similar nt/AA in earlier epidemics.

### Signature substitutions between lineages with successful and limited local transmission

In an attempt to unravel the molecular basis of the ON1 lineages in Kilifi with varied local transmission outcomes, we performed a similar genome-wide comparative scan between the consensus of genomes of viruses with successful (lineages LI and LII) and those with limited local transmission (lineage LIII) for characteristic signature polymorphisms. We identified 33 SNPs between these two groups of lineages, *supplementary sheet 3*, of which nine resulted in non-synonymous changes; five in G, two in F and one each in M2-2 and L. In three of these nine SNPs with non-synonymous substitutions between the two groups of lineages, lineage LIII possessed similar nucleotides as the GA2 viruses (G: codons P274L and P310L, and F: codon A122T). Whether these polymorphisms that characterize the two groups of lineages are neutral mutations or influence local transmission of the virus warrants further investigation.

### Patterns of selective pressure

It is expected that different codon sites in genes would be under differential evolutionary pressures. We conducted selection analysis on all 11 RSV ORFs for the dataset. ORF-wide episodic diversifying selection was only detected in the NS1 and M proteins. A total of nine positively selected codon sites were identified within the G (73, 201, 250, 251, 273, 310), NS2 (15) and the L (2030, 2122) by at least one method. Notably, sites 273 and 310 within the G protein detected to be under positive selection also had the signature SNPs described above between ON1 and GA2 viruses. However, the number of positively selected sites could have been underestimated in the analysis that was limited to Kilifi RSV-A genomes and care should be taken while interpreting these results as some of the positively selected sites were only detected by one method and at default (less stringent) cut-offs.

## Discussion

Here we report an in-depth analysis of local and global RSV genotype ON1 evolution and transmission using whole genome sequence data. We describe RSV-A genomic diversity and identify polymorphisms with the most potential in influencing RSV evolution and phenotype. Utilizing genomes from samples collected between 2010–2016, including 184 complete genomes from Kilifi alone, we obtained a finer resolution on the pattern of RSV introductions, persistence and evolution in a defined location, and the changes within the genome that might be important for the survival of the virus.

Genetic variation not only provides important insights into RSV relatedness by which to infer transmission but also highlights potential functional changes in the virus. From our analysis, we find that substitutions are widespread across the RSV genome but occur at higher frequency within the structural proteins (G and F) and in parts of the polymerase (L). The G protein has the most genetic flexibility of the RSV ORFs to accommodate frequent substitutions including large duplications, and previous studies have described epitope positions within this protein that characterize escape mutants selected by specific monoclonal antibodies or by natural isolates (5,68,70,71). Site p27 in the F protein with two signature substitutions has been shown to possess greater binding activity in sera from young children (<2 years) than any of the other antigenic sites in the F protein and may be responsible for group specific immunity due to its great variability between RSV-A and RSV-B viruses (72). However, the implications of the substitutions in the L protein of the ON1 viruses remain unclear, but considering both its role in genome replication and the presence of the 72-nucleotide duplication in the G ORF, we posit that either (i) these polymorphisms might have resulted in a sloppy polymerase, that further resulted in a ^1^ slip that generated the 72-nucleotide duplication in the G ORF (73), or (ii) the 72-nucleotide duplication in the G presented a larger metabolic challenge in replicating a larger genome and thereby facilitating adaptive polymorphisms within the polymerase (74). While we also found a considerable number of SNPs in other ORFs other than the G, F and L proteins, only a very minor proportion of those changes resulted in amino acid substitutions implying very strong purifying selection in these portions of the genome.

Based on distinct phylogenetic clustering of ON1 and GA2 viruses in five ORFs, the emergence of ON1 may be characterized by additional substitutions across the genome in addition to the 72-nucleotide duplication within the G gene. However, assuming ON1 diverged from GA2 and through a single ancestral virus, it is unclear whether the multiple signature substitutions differentiating ON1 from GA2 viruses all: arose from that single split event or have been acquired progressively over time. In case of the latter, it is unclear the chronology of the changes across the different ORFs. Understanding how and which mutations define the emergence of a new RSV variant may be important in describing substitutions that are either crucial for the survival of the variant and/or of some complementary structural or functional integrity. It is also likely that some of these substitutions are nothing more than genetic hitchhikers. Notwithstanding this lack of clarity on ON1 emergence, it has been shown for influenza A viruses that linked selection amongst antigenic and non-antigenic genes influences the evolutionary dynamics of novel antigenic variants (75). Further, it has been demonstrated experimentally that adaptive evolution is a multi-step process that occurs in waves (76). There is an initial adaptive wave that occurs rapidly and is characterized by founder or gatekeeper mutations. Thereafter, additional waves of evolutionary fine-tuning occur (77). Similar studies in RSV would be important in determining if such dynamics do characterize their evolutionary history and also might inform the design of an RSV vaccine.

Undoubtedly, ON1 is rapidly replacing GA2 in Kilifi suggesting that this variant has some fitness advantage. We previously showed that genotype ON1 viruses did not result in more severe disease compared to GA2 viruses in Kilifi (32). However, there are conflicting reports globally with some indicating that ON1 is more virulent and others reporting ON1 being less virulent than GA2 (78,79). Even with the discordant results, which may also be due to differences in study populations and analysis methods, there might be phenotypic differences between viruses belonging to these two genotypes. Identification of such phenotypic differences and the potential drivers might augment our current understanding of the pathogenesis of this virus.

Observations from this study using whole genomes reinforce previous findings based on partial G-gene sequences (17,18,22,32) that RSV epidemics are characterized by the introduction and circulation of multiple variants. In addition, persistence within the community seems to be sustained by only a proportion of these introductions. We have characterized genomic substitutions that distinguish between successful and dead-end ON1 lineages in Kilifi. Nonetheless, it is evident that besides viral genetic factors there could be other determinants of successful onward transmission of a virus lineage considering that the non-persistent ON1 strains in Kilifi were abundant in other parts of the world albeit with varied frequencies relative to other genotypes. Such determinants warrant further investigations and could include the host factors (e.g. births, immunity, genetics, contact patterns and mobility) and environmental factors (e.g. temperature, rainfall and humidity).

We live in times of rapid global movement of people, which may influence the spread of infectious diseases. The observation that_most of the Kilifi sequences clustered with sequences from Europe and Asia suggests that RSV introductions into Kilifi originate predominantly from these two continents. It might not be surprising that Europe is a primary source of RSV introduction into Kilifi, or even a destination for viruses from Kilifi, considering that it accounts for the largest single group of tourists to Kenya (80). In addition, the increasing Chinese economic interests in Africa (including Kenya) in becoming Africa’s largest trade partner has resulted in an influx of Chinese into Africa for trade, work and tourism (81). However, there are far too few partial ON1 sequences from Africa (only from Kenya, South Africa and Nigeria) and no ON1 genomes from outside Kilifi Kenya to help define intra-African spread dynamics in detail (which we hypothesize might be more impactful on the many local introductions). In fact, a recent study suggests that in the recent past domestic tourism accounts for more than half of the growth in Kenya’s tourism (82). As such, availability of sequences from across the country would be critical in deciphering if and how such tourist activities influence virus transmission patterns in Kenya. Accordingly, such observations could be helpful in the design of future RSV transmission intervention strategies.

## Supporting Information

**S1 Fig:**
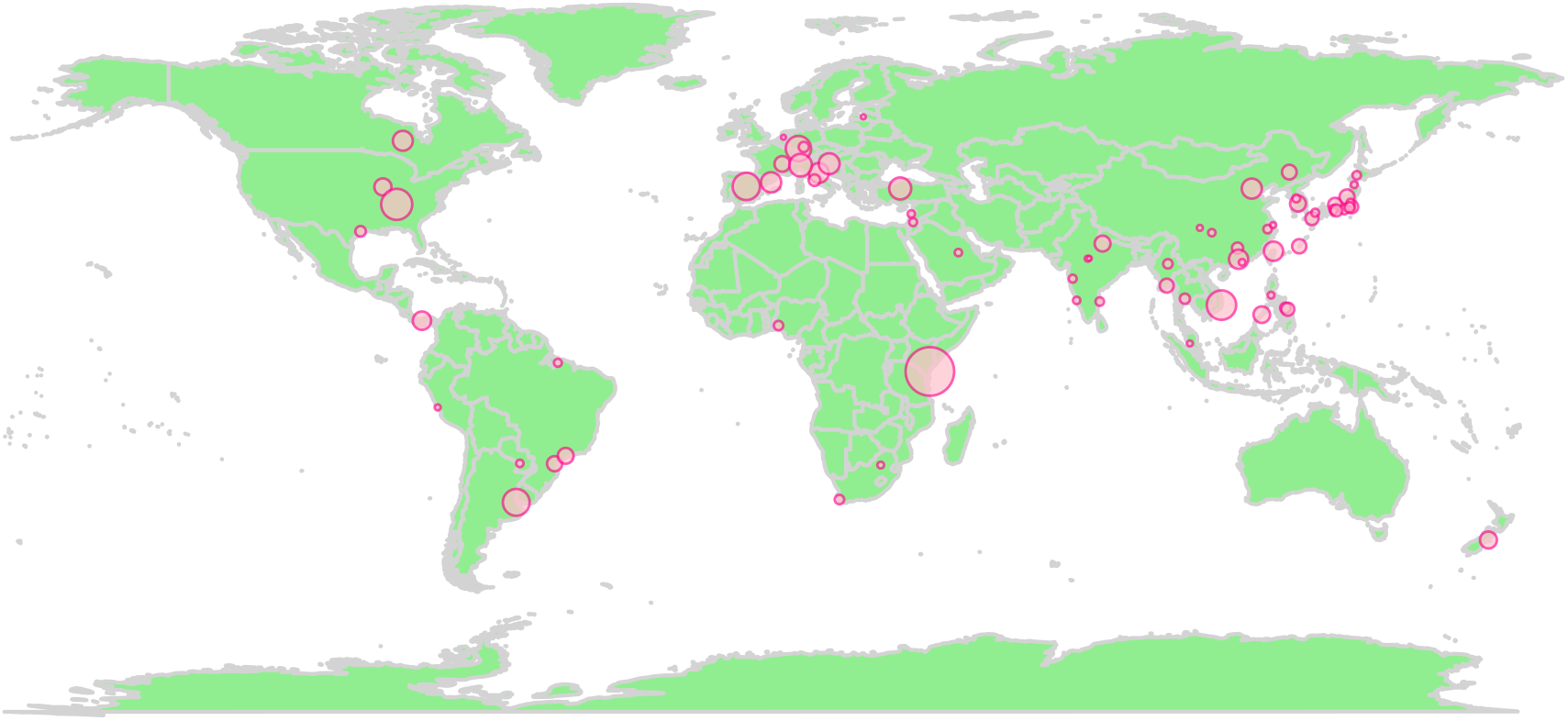
Sampling locations of the global ON1 G-gene dataset. A map showing source country locations of the global ON1 G-gene sequences dataset analyzed here with circles representative of relative proportion of contributing sequences by country

**S2 Fig:**
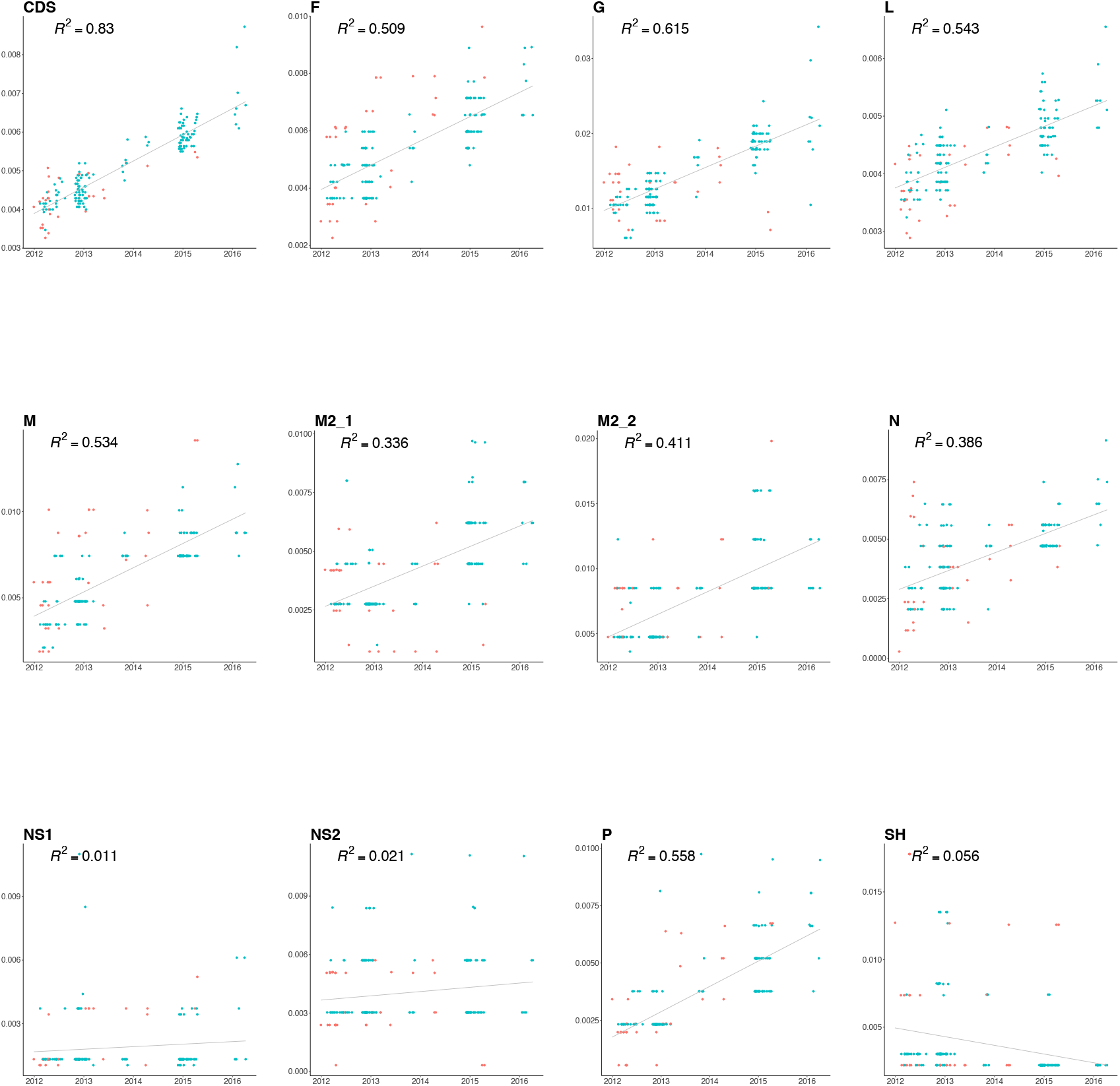
Root-to-tip regression analysis of Kilifi RSV-A ORFs. A root-to-tip regression analysis of ML trees from whole genomes and 11 separate coding regions, with the points colour coded by genotype; GA2 (red) and ON1 (cyan).

**S3 Fig:**
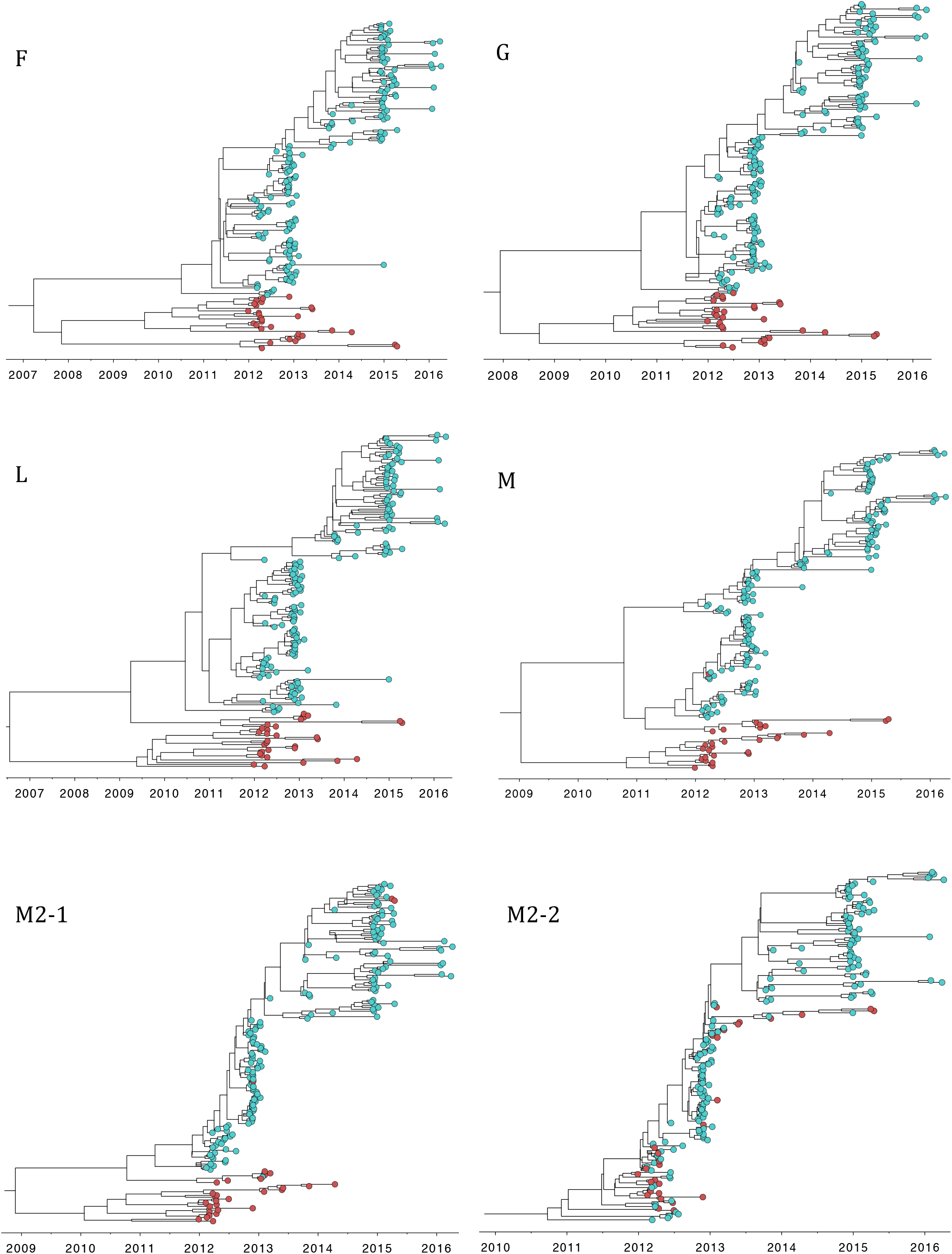
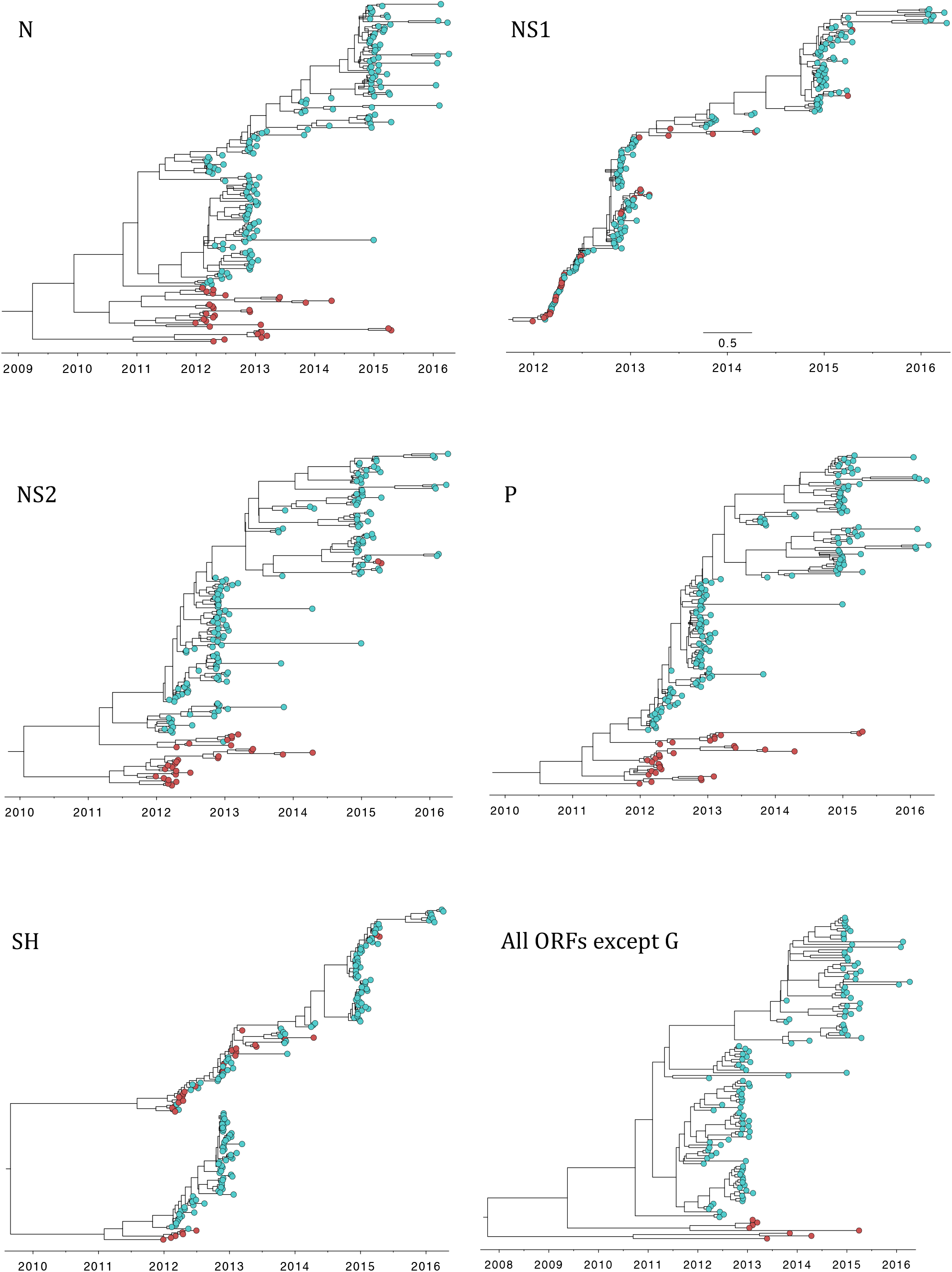
BEAST MCC ON1-GA2 divergence trees for different ORFs. MCC trees inferred from different ORFs of 184 RSV-A complete genome sequences from Kilifi with the tips colour coded by genotype, i.e. ON1 (cyan) and GA2 (red).

**S1 Table: Model selection to infer time-structured phylogenies**

**S2 Table: Study samples and genomes details**

**S3 Table: SNPs identified from dataset of all Kilifi genomes**

**S4 Table: Signature SNPs between ON1 and GA2 viruses**

**S5 Table: Signature SNPs between successful and limited transmission ON1 lineages**

